# SliceIt: A genome-wide resource and visualization tool to design CRISPR/Cas9 screens for editing protein-RNA interaction sites in the human genome

**DOI:** 10.1101/654640

**Authors:** Sasank Vemuri, Rajneesh Srivastava, Quoseena Mir, Seyedsasan Hashemikhabir, X. Charlie Dong, Sarath Chandra Janga

**Author notes:** Both the authors contributed equally to this study and should be considered as joint first authors (Sasank Vemuri, Rajneesh Srivastava). Correspondence should be addressed to: Sarath Chandra Janga, 719 Indiana Avenue Ste 319, Walker Plaza Building, Indianapolis, Indiana – 46202.

## Abstract

Several protein-RNA cross linking protocols have been established in recent years to delineate the molecular interaction of an RNA Binding Protein (RBP) and its target RNAs. However, functional dissection of the role of the RBP binding sites in modulating the post-transcriptional fate of the target RNA remains challenging. CRISPR/Cas9 genome editing system is being commonly employed to perturb both coding and noncoding regions in the genome. With the advancements in genome-scale CRISPR/Cas9 screens, it is now possible to not only perturb specific binding sites but also probe the global impact of protein-RNA interaction sites across cell types. Here, we present SliceIt (http://sliceit.soic.iupui.edu/), a database of in silico sgRNA (single guide RNA) library to facilitate conducting such high throughput screens. SliceIt comprises of ~4.8 million unique sgRNAs with an estimated range of 2–8 sgRNAs designed per RBP binding site, for eCLIP experiments of >100 RBPs in HepG2 and K562 cell lines from the ENCODE project. SliceIt provides a user friendly environment, developed using advanced search engine framework, Elasticsearch. It is available in both table and genome browser views facilitating the easy navigation of RBP binding sites, designed sgRNAs, exon expression levels across 53 human tissues along with prevalence of SNPs and GWAS hits on binding sites. Exon expression profiles enable examination of locus specific changes proximal to the binding sites. Users can also upload custom tracks of various file formats directly onto genome browser, to navigate additional genomic features in the genome and compare with other types of omics profiles. All the binding site-centric information is dynamically accessible via “search by gene”, “search by coordinates” and “search by RBP” options and readily available to download. Validation of the sgRNA library in SliceIt was performed by selecting RBP binding sites in Lipt1 gene and designing sgRNAs. Effect of CRISPR/Cas9 perturbations on the selected binding sites in HepG2 cell line, was confirmed based on altered proximal exon expression levels using qPCR, further supporting the utility of the resource to design experiments for perturbing protein-RNA interaction networks. Thus, SliceIt provides a one-stop repertoire of guide RNA library to perturb RBP binding sites, along with several layers of functional information to design both low and high throughput CRISPR/Cas9 screens, for studying the phenotypes and diseases associated with RBP binding sites.

## 1. Introduction

CRISPR (Clustered Regularly Interspaced Short Palindromic Repeats) is identified as a defense system that protects bacteria and archaea from mobile genetic elements [1–3]. This RNA guided interference mechanism has been successfully employed in eukaryotic cells (both in vitro and in vivo) by Zhang [4] and Charpentier [5] groups. It is now established that sgRNAs (single guide RNAs) can be engineered to target a 17-20 bp stretch of DNA sequence preceding a protospacer adjacent motif (PAM) [6]. CRISPR/Cas9 system has been crucial in multiple disciplines but is especially useful for understanding the gene function by manipulating precise genomic locations [6–8]. This emerging technology enables the re-investigation of multilayered functional dependency of regulating molecules such as kinases, transcription factors (TFs), long non-coding RNAs (lncRNAs), and other protein coding gene groups, to provide specific perturbations and to study their comprehensive genome wide effects [7, 9–12].

RNA binding proteins (RBPs) post-transcriptionally regulate a variety of biological processes including 5’ capping, 3’ polyadenylation, polyadenylation, splicing, RNA editing, transport, localization, stability, degradation and translation [13–16]. These evolutionary conserved proteins [15] are often implicated in modulating the post-transcriptional networks in many human diseases including cancer [17, 18] and neurodegenerative disorders [14, 19]. Several crosslinking and immunoprecipitation (CLIP)-seq protocols have been developed over the years [20–22] to delineate the molecular interaction of RNA binding proteins (RBPs) and their target RNAs at single-nucleotide resolution in a cell. However, most of the millions of binding sites identified from these high throughout CLIP-seq studies do not have functional evidence for their contribution to the fate of the RNA molecule, except perhaps binding the RNA target, creating an ambiguity with little or no functional relevance of these interactions in cellular context [23, 24]. It has also been argued that various CLIP-seq protocols don’t agree with each other in recovering binding sites and often produce noisy signals resulting in a number of false positive binding sites [25].

With the advent of genome modification via CRISPR/Cas9 systems [6–8], it is possible to investigate the functional connectivity between RBPs and RNAs, specifically with high resolution. CRISPR/Cas9 system can potentially be employed to investigate the functional aspect of localized RBP-RNA interactions in cells (Figure 1). This system, though originally developed to directly target and edit DNA, recent studies also report the use of variants of Cas9 for tracking RNA [26, 27]. However, the system’s efficiency to access/edit RNA is highly compromised with low signal to noise ratio and accuracy. In addition, editing RNA (which can’t be repaired by cellular pathways) will not enable the use of expression of the target RNA molecule as a proxy to measure the functionality of the binding site. CRISPR/Cas9 system is precise and cheaper compared to other gene editing techniques. Apart from single target site perturbation, it can also be used to target multiple loci simultaneously with different sgRNAs and using a single Cas9 variant. The ability of using single Cas9 protein with multiple sgRNAs opens the doors for high throughput editing of target loci. Therefore, to understand the impact of RBP binding sites perturbation in human cells, we propose to use CRISPR/Cas9 system to edit the DNA locations where RBP binds to RNA.

**Figure 1.**
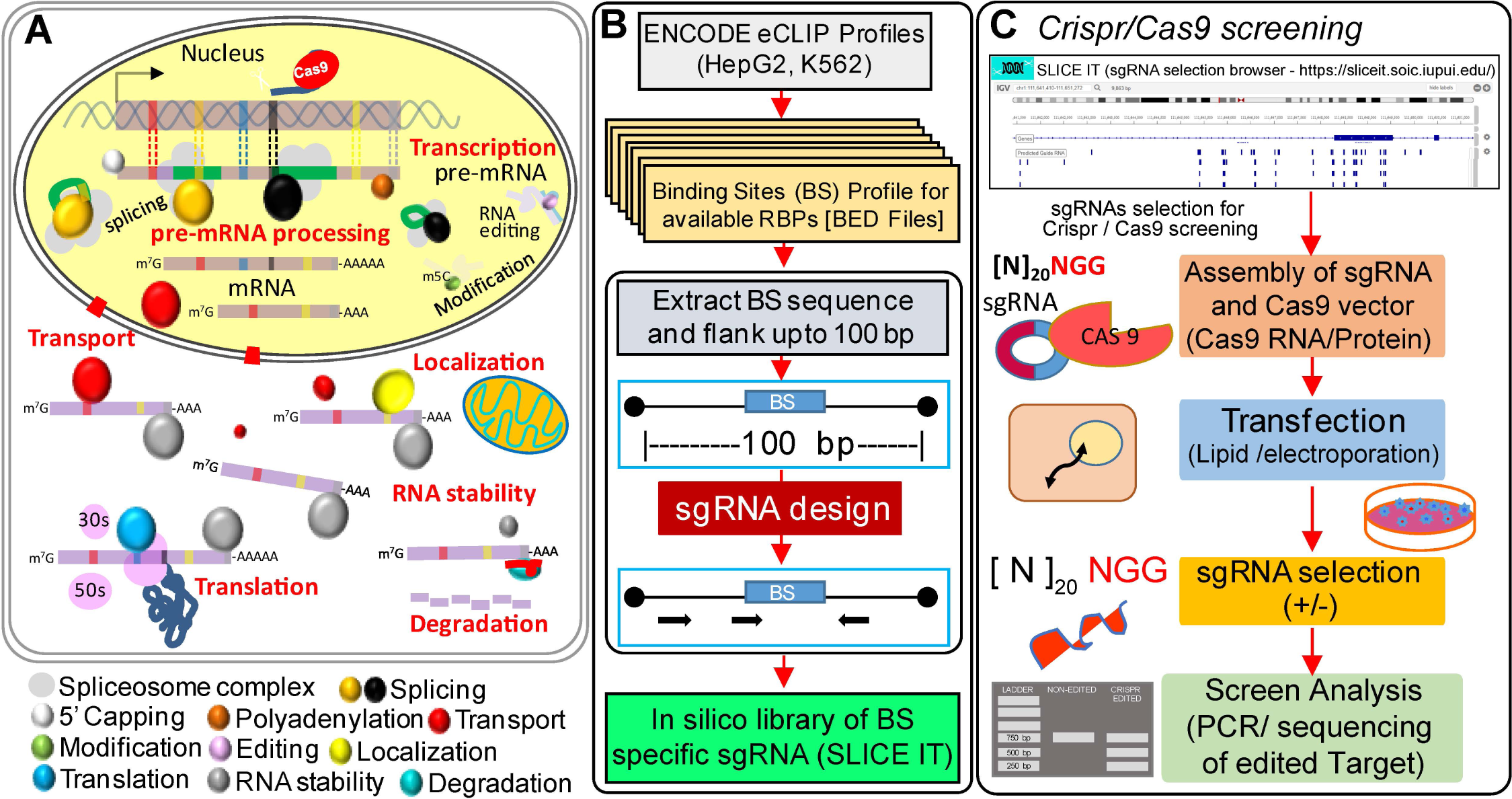
Strategy for validation of RBPs’ binding sites. (A) A hypothesis driven CRISPR/Cas9 model for determining the effect of sgRNA on an exon proximal to targeted binding site (B) Construction of SliceIt, an in-silico guide RNA library for RBPs’ binding sites (C) Proposed approach for designing CRISPR/Cas9 screening experiments based on the compendium of sgRNAs from SliceIt, for perturbation of binding sites.

SliceIt is the first comprehensive database of in silico sgRNA library to edit the currently known millions of protein-RNA interaction sites in the human genome. It stands as a one-stop portal for designing CRISPR/Cas9 screens for functional dissection of post-transcriptional regulatory networks. SliceIt enables the designing of multiple sgRNAs for each binding site based on user’s desired location for editing the genome by defining specificity and efficiency thresholds, to facilitate uncovering their functional role in modulating post-transcriptional regulation of a transcript. Predicted sgRNAs are available to be visualized in a genome browser with additional layers of information such as SNPs and cis-exon expression across human tissue types. SliceIt also provides an option to download data for every search query in CSV or Excel file format. SliceIt uses Flask micro framework in the backend to efficiently parse results and to communicate with user interface and the databases. In order to handle multiple queries efficiently, dynamic implementation of multiprocessing is used to parallelize querying process from Elasticsearch cluster to effectively reduce query search time by several fold.

## 2. Materials and Methods

### 2.1. Data collection and processing

We obtained eCLIP [28] based binding profiles of scores of RBPs in (hepatocellular carcinoma (HepG2) and chronic myelogenous leukemia (K562) cell lines from the ENCODE project [29]. In total, our dataset comprised of 2.23 million unique binding sites for 68 RBPs in HepG2 and 2.38 million unique binding sites for 86 RBPs in K562 cell line. We downloaded the genomic coordinates of these RBPs’ eCLIP profiles (in .bed file format), parsed and used them for prediction of sgRNAs localized to each binding site of RBPs. In addition to CLIP profiles, we also downloaded dbSNPs [30], GWAS catalog [31] and exon expression profiles for 53 human tissues from the GTEx project [32] (GTEx data was extracted by using recount workflow [33, 34]) and integrated with SliceIt, to provide comprehensive information for a genomic region of interest.

### 2.2. Prediction of sgRNAs around RBPs’ binding sites

A typical sgRNA comprises of a 19-20 base long oligonucleotide that could be designed to target user defined homologous sequence on the host genome. In this study, we used CRISPR-DO [6] to design sgRNAs targeting all possible RBP binding sites from various CLIP experiments [28] in HepG2 and K562 cell lines from ENCODE [29]. CRISPR-DO requires an input genomic region in bed format along with other essential metrics such as genome assembly and spacer length. We customized the RBP binding site coordinates obtained from the ENCODE project for human reference genome (hg38) and employed a flanking distance of ±50bp from the mid-point of the binding site (if BS length is <100 bp) and automated the standalone CRISPR-DO software (http://cistrome.org/crispr/source) to predict potential sgRNAs per binding site using ad-hoc scripts. This enabled the development of an in-house pipeline for sgRNA design to millions of RBP binding sites on a compute cluster.

### 2.3. Processing of CRISPR DO outputs

CRISPR-DO provides genomic location, 30 bp sequence (i.e. 20 bp sgRNA sequence + PAM + 7 bp flank sequence), sgRNA strand orientation, specificity score, efficiency score and other flagged annotations for all possible sgRNA predictions per genomic region of interest. Predicted sgRNAs and other information for each binding site were tagged with respective queried binding site (BS) coordinates and corresponding cell line as well as RBP information. These predictions were concatenated into a single file. We also computed the distance between the mid-point of the BS and PAM site, to include it as an additional column in the processed outputs.

### 2.4. Database construction and implementation

SliceIt is a database implemented in elasticsearch with a web interface that was developed using Bootstrap 4.0 software. The database is hosted at https://sliceit.soic.iupui.edu/. SliceIt currently allows users to search by gene name or Ensembl ID, coordinates and by RNA binding protein name at various efficiency and specificity thresholds.

#### 2.4.1. Data Interface

SliceIt interface comprises of three different search options to query the data and the search result of all of these primary functions include (i) Retrieval of information on guide RNAs corresponding to the binding sites (ii) Retrieval of SNPs and GWAS information that fall within the region of interest (iii) exon expression levels across tissues that are in 500 bp proximity to region of interest and are defined as cis-exons in this study (iv) Visualization of data in IGV JS genome browser. When a user queries for a gene, SliceIt provides a list of various genes and their corresponding Ensembl IDs in the form of an auto suggest dropdown box. For each search the recommended and default cut-off for efficiency and specificity scores for selection of sgRNAs are 0.3 and 50 respectively. These numbers are recommended for selection of optimal guide RNA design by CRISPR-DO tool. Results retrieved in SliceIt are organized into the following sections

a. Annotations: The output page displays annotation information obtained from Ensembl for search based on gene name or coordinate range within a chromosome. These annotations cover the gene name, description, location coordinates, strand information along with a link to Ensembl database to access more information.
b. Genome Browser: SliceIt helps visualize the locations of binding sites and sgRNAs with the help of IGV JS genome browser embedded in the output page. By default, the tracks that are displayed include (i) hg38 reference genome (ii) GWAS (iii) SNP from dbSNP (iv) Binding sites in HepG2 cell line (v) Binding sites in K562 cell line (vi) Predicted sgRNAs. The users have an option to remove and add any of these tracks, change track color, name and height. Apart from the default tracks that are loaded for every search query, users can also load various other tracks by using drop down menus. These options for additional tracks include exon expression tracks for various tissues, HepG2 and K562 binding sites tracks for individual RBPs. SliceIt also has an additional functionality that allows users to add their own data in the form of a track in genome browser by using the “Add custom track from URL” section on the results page. This allows users to add indexed BED, BAM, Wig, Bigwig and BedGraph file formats that are hosted on an external server.
c. Data Tables: The retrieved raw data is displayed in “Data View” tab in a tabular form with various options such as CSV and Excel export, search filter by coordinates, efficiency or specificity and column sorting. For each binding site, SliceIt provides 5 different sgRNAs that are filtered based on the highest efficiency and specificity scores. The data view also has 3 other sub-tabs that provide data regarding SNPs, GWAS and Cis-exon expression.

#### 2.4.2. Backend

In the backend, SliceIt runs on Python’s Flask micro web framework to efficiently process the query, parse the data and return output. Predicted sgRNAs, dbSNPs, GWAS and Exon expression data is stored in elasticsearch (https://www.elastic.co/) that is hosted on an external cluster. For each search, the query input is passed to the backend via flask framework and SliceIt automatically designs various queries to efficiently retrieve data from elasticsearch and Ensembl. This process comprises of 4 components

a. Generating annotations: Annotation data is retrieved from Ensembl database with the help of API.
b. Generating elasticsearch queries: SliceIt is designed with flexibility in mind and hence the complete code is modular and scalable. As the user submits a search query, the backend passes the query information to various modules that automatically design specific elasticsearch queries to retrieve data for binding sites, SNPs, GWAS and exon expression. This dynamic workflow is implemented to retrieve data that is in closest proximity by distance in base pairs to the binding sites. This is made possible by incorporating painless groovy-style scripting language within each json query object. The painless scripting extends Java’s programming functionality within a search query and this allows fast, safe and very specialized retrieval of data for each search input which is not feasible with any mySQL or relational databases.
c. POST queries to elasticsearch: SliceIt uses a fundamental component from dynamic programming to efficiently process the search queries. Each search is broken down into sub-problems and an optimal substructure is developed for query processing. Once the queries are generated, multiprocessing is implemented to parallelize each query to reduce data retrieval speed by up to 6-fold. As elasticsearch uses RESTful API, the queries that are generated are in JSON format. Each parallel process sends the JSON query as “POST” request to elasticsearch cluster and then output is returned in the form of JSON objects. Each output JSON object is independently collected by multiprocessing pool for further parsing.
d. Output parsing and template rendering: The query output is parsed to obtain relevant data and passed to the frontend to be displayed in various tabs via bootstrap front-end.

This pipeline takes only few seconds to process and render an output. While most other alternative tools run an algorithm in the cloud to generate guide predictions, SliceIt takes advantage of precomputed data to achieve a speed that is several folds faster than other comparable tools. The database is free for all users and the complete dataset can be provided upon request.

#### 2.5.1 sgRNA design for experimental validation

SliceIt serves as the first comprehensive predictive engine for designing sgRNAs to edit the currently known millions of protein-RNA interaction sites in the human genome that augment to conduct high-resolution binding site block/silencing experiments. We designed 2 sgRNAs (5’- CTGGTAGGGGAGTCAAGAGA-3’ for BS chr2:99157353-99157403 and 5’- AGTATGGATTAAATAAAGGA-3’ for BS chr2:99159478-99159514) in LIPT1 (Lipoyltransferase 1) gene locus from SliceIt.

#### 2.5.2 CRISPR Cas9 system

Lentiviral vector digestion, oligo annealing and cloning into digested vector: Lentiviral CRISPR v2 plasmid (a gift from Feng Zhang (Addgene plasmid # 52961; http://n2t.net/addgene:52961; RRID:Addgene_52961) was digested with BsmBI and dephosphorylated with alkaline phosphatae for 2 hrs at 37°C. Digested plasmid was purified from gel using QIAquick gel extraction kit as per manufacturer’s instructions. Oligos were phosphorylated and annealed using T4 PNK (NEB M0201S) enzyme and T4ligation buffer (ATP added) in a thermocycler using following parameters: 37°C for 30min, 95°C for 5 min and ramp down to 25°C at 5°C/min. Annealed oligos are diluted at 1:200 dilution into sterile water. Diluted oligos were ligated with digested plasmid using T4DNA ligase, incubated over night at 16°C. Lentiviral plasmid was transfected into Stbl3 bacteria (Invitrogen C7373-03) using heat shock transfection method. For each oligo, three clones were selected for plasmid isolation. Plasmids were isolated using GeneJET Plasmid Miniprep Kit (K0503) and sent for sanger sequencing.

#### 2.5.3 Lentiviral packaging

One day before transfection 2.5×10^5^ HEK293T cells were plated (6 well plate) in DMEM supplemented with 10% heat-inactivated fetal bovine serum (FBS). Cells were incubated at 37°C overnight to get a confluency of 70%. Transfection was carried out using polyethyleneimine (PEI) method with the ratio of PEI:pTarget:pVSVg:RRE:REV; 16:3:1: 2:2. In a sterile tube, total 3ug of DNA following the ratios was diluted to 200ul of serum-free DMEM. PEI (2ug/ul) based on a 2:1 ratio of PEI(ug): total DNA(ug) was added to diluted DNA. Mix was incubated for 15 min at room temperature. For each binding site, one oligo was transfected individually into the cells, to purturb the binding site. After 72 hrs, lentiviral particles were harvested and concentrated at 3,000g for 5min at 4°C. The supernatant is filtered through a 0.45um filtration on ice using synringe filter. HepG2 cells were freshly cultured in 6 well plate for 24hrs, followed by transduction using lentiviral concentrate. Cells were incubated for one week followed by which puromycin treatment was given for positive selection of transduced cells. GFP and plasmid insert was used as positive control. Fluorescence microscope was used to check transfection and transduction efficiency. After 1 week puromycin concentration was reduced, since the cell number was low. Cells were split after 50% confluency of wells. After 1 week of incubation, cells were harvested in two tubes. From one aliquot, RNA was isolated using Tri-reagent, cDNA generated and real-time PCR was run for analyzing the expression level of proximal exons. In order to validate gene modifications, DNA was isolated from the second aliquot and sent for sanger sequencing.

## 3. Results and Discussion

RBP driven post-transcriptional regulation likely depends on its binding efficiency to its target location [35, 36] (Figure 1A). This phenomenon is highly crucial for several key biological processes including in development [37–40] and differentiation [13, 41–46]. It can be studied by measuring the expression level of the target RNA or proximal/neighboring exon to that of the binding site of interest. We hypothesize that, perturbation of the RBPs’ binding sites or its equivalent position on DNA can potentially promote dysregulated function of the post-transcriptional target RNA molecule, enabling the functional dissection of the millions of binding sites of RBPs being discovered by CLIP [47] and related technologies [20–22]. Cas9 system has been extensively utilized to edit the genomic loci of interest [6, 48]. However, it has not been used to systematically understand the impact of RBP binding site perturbation in human cell types. Thus, we employed this “cause” and “effect” model of regulation by perturbing equivalent binding site on DNA using Cas9 system, where dysregulation phenotype can be measured by expression analysis of the target RNA feature, such as inclusion or exclusion of exon. Briefly, we downloaded the BS profiles for available RBPs from the ENCODE project and preformatted the BS regions by flanking them up to 100 bp. We employed CRISPR-DO to design sgRNAs for each binding site. All predicted sgRNAs were deposited into a database called SliceIt (see Figure 1B and Materials and Methods section). SliceIt facilitates designing CRISPR/Cas9 screens in both low (as illustrated in Figure 1C) and high throughput modes, by enabling parametric flexibility for designing sgRNAs along with the ability to filter the binding sites for the presence of SNPs, GWAS hits and exon expression alterations across a wide range of human tissue types.

### 3.1. Overview of SliceIt

In this study, we present SliceIt (https://sliceit.soic.iupui.edu/), a database and visualization tool providing a comprehensive summary of in silico sgRNA (single guide RNA) library, to facilitate rational design of CRISPR/Cas9 experiments in both low and high throughput fashion to perturb the protein-RNA interaction sites. We used CRISPR-DO [6] to design ~4.9 million unique sgRNAs targeting all possible RBP binding sites resulting from eCLIP experiments of scores of RBPs in HepG2 and K562 cell lines (from ENCODE) (see Materials and Methods). SliceIt provides a user-friendly environment, developed in highly advanced search engine framework called Elasticsearch. It is available in both table and genome browser views facilitating the easy navigation of RBP binding sites, sgRNAs, SNPs and GWAS hits, while querying for a gene, RBP or region of interest. It also provides exon expression profiles across 53 human tissues from the GTEx project (https://gtexportal.org/home/) [32], to examine locus specific expression changes proximal to the binding sites, to enable rational design of experiments in specific tissue/cell types. Users can also upload custom tracks in various file formats (in browser) to navigate additional genomic features in hg38 human genomic build. This custom track upload and navigation feature, in addition to the datasets already integrated into SliceIt, provide a functional context for user-generated datasets. All the binding site centric information is dynamically accessible via “search by gene”, “search by coordinate” and “search by RBP” and readily available to download. SliceIt is the first comprehensive predictive engine for designing sgRNAs to edit the currently known millions of protein-RNA interaction sites in the human genome. It is a one-stop repertoire of guide RNA library and RBP binding sites along with several layers of functional information, to design high throughput CRISPR cas9 screens for studying the phenotypes and diseases associated with the binding sites of RBPs and to functionally dissect the post transcriptional regulatory networks.

### 3.2 Characteristics of sgRNA repertoire available in SliceIt

We downloaded ~4.6 million binding sites corresponding to 108 RBPs across the two cell lines (2.23 million unique sites in HepG2 and 2.38 million unique sites in K562) from the ENCODE project [49] (Figure 2A). These binding sites were flanked to 100 bp (if <100 bp) and queried for sgRNA prediction using CRISPR-DO (see Materials and Methods). This resulted in a repository of ~4.9 million unique sgRNAs (3.73 million and 3.04 million unique sgRNAs for HepG2 and K562 cell lines respectively) predicted for ~4.6 million binding sites. Binding sites for each RBP and corresponding unique sgRNAs were log10 transformed and illustrated as a heatmap in Figure 2A. We also calculated the ratio of number of sgRNAs and binding sites for each RBP, representing an estimate of the average number of sgRNAs designed per binding site for each RBP in both the cell lines (Figure 2A). SliceIt comprises of a relatively unbiased collection of sgRNAs (2-8 sgRNAs per binding site) for the RBPs included in the database, making it an easily accessible and user-friendly web interface for designing the targeted post-transcriptional dysregulation experiments using Cas9 system. Additionally, we observed that the total distribution of designed sgRNAs decreased with increasing efficiency as well as specificity (Figure 2B). Several CRISPR based experiments have shown that Cas9 directed double strand breaks occur mostly in close proximity to PAM region of targeted genomic loci [6]. Thus, we investigated the positional occurrence of designed sgRNAs by calculating the distance of binding site from PAM. We observed that most of the designed sgRNAs were proximal to the midpoint of the binding sites with an exponential decline as distance increased from the center of the binding site (Figure 2C). It is noteworthy to mention that the total number of designed sgRNAs and made available via SliceIt varied between chromosomes and were generally correlated with the size of the chromosome (Figure 2D).

**Figure 2.**
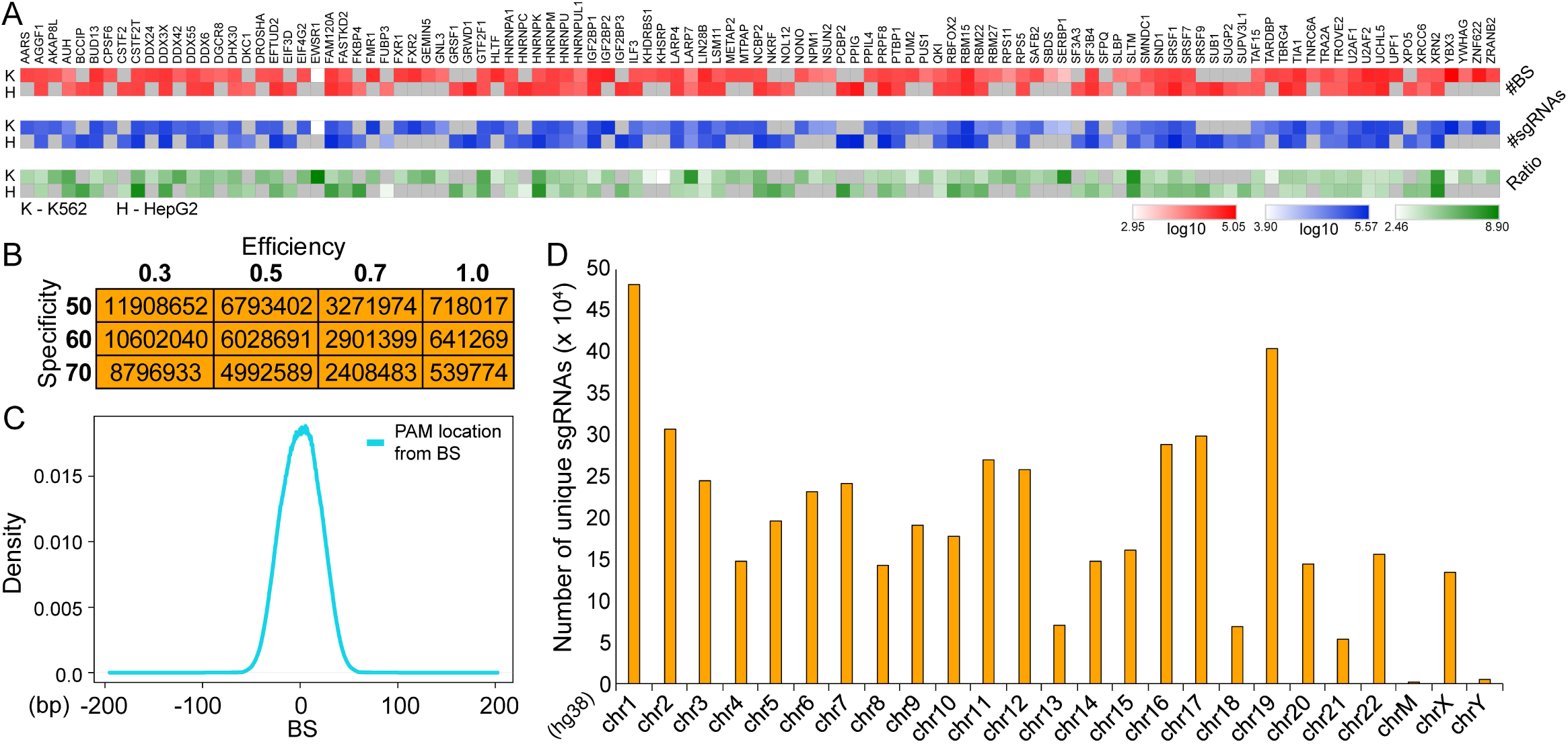
**(A)** Heatmap showing the number of unique binding sites (log10 transformed -red) and sgRNAs (log10 transformed-blue) and the ratio of the number of sgRNAs and binding sites (green) for each RBP, representing an estimated average sgRNAs designed per binding site for each RBP in both the cell lines (H-HepG2 and K-K562). RBPs for which currently no binding site information is available from ENCODE project for either cell line are greyed out. **(B)** Distribution of the total number of designed sgRNAs available from SliceIt as a function of the predicted efficiency and specificity scores. **(C)** Density plot showing the distribution of distances between sgRNA’s PAM location and the mid-point of the targeted binding site, for all the sgRNAs available from SliceIt. **(D)** Distribution of the absolute number of sgRNAs across human chromosomes present in SliceIt.

### 3.3 SliceIt database construction, visualization and accessibility

SliceIt is an efficient search engine, consisting of several layers of omics data including RBP binding site profiles, SNPs, GWAS and tissue-specific exon expression levels (GTEx) under the niche of Flask server module. The complete pipeline as illustrated in Figure 3A, details on how the flask server interacts with front-end, retrieves and parses data from elasticsearch for providing an output. Basically, this pipeline takes only a few seconds to search the query, process and render an output. While most other alternative tools run an algorithm in the cloud to generate guide RNA predictions, SliceIt takes advantage of precomputed data to achieve a speed that is several fold faster than other comparable tools. For detailed structure of the resource and how this pipeline functions, refer Data Interface (2.4.1) and Backend (2.4.2) in Materials and Methods section. SliceIt enables searching a transcribed region with RBP binding sites in the genome for designed sgRNAs. This is facilitated by allowing the user to search for gene region, genomic co-ordinates and for targets of an RBP (Figure 3B). For instance, a user could search for the sgRNAs that can be designed to edit the binding sites of a member of the RBFOX family of RBPs or search for a specific genomic region defined by chromosomal co-ordinates. Alternatively, a user can visualize the target binding sites of a selected RBP for which sgRNAs satisfy specific design thresholds. For each search query, SliceIt outputs query annotations, data visualization with IGV JS genome browser, SNPs in the region, GWAS SNPs in the region, predicted sgRNA and cis-exon expression information in tabular form as shown in Figure 3C. Cis-exon expression information across human tissues obtained from the GTEx project is displayed in Fragments Per Kilobase of transcript per Million mapped reads (FPKM) units and is color coded to indicate various ranges of expression levels (see Materials and Methods). An expression level of less than 1 FPKM is displayed in dark red, greater than 1 and less than 10 FPKMs in yellow, greater than 10 and less than 100 FPKMs in orange and an expression level higher than 100 FPKMs is displayed in green.

**Figure 3.**
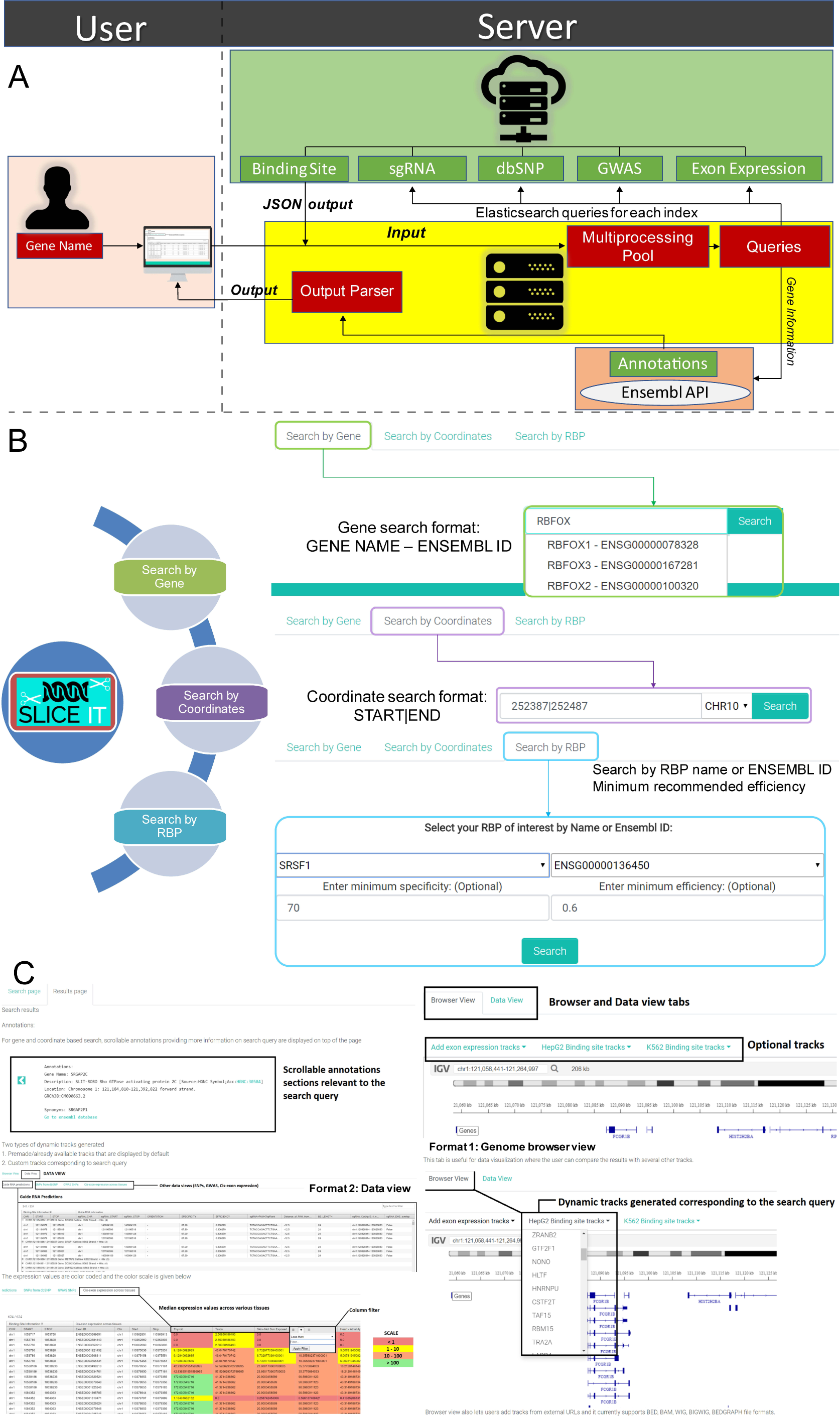
**(A)** Search processing pipeline showing the functionality of SliceIt’s communication with frontend and elasticsearch cluster to parse and display query output. **(B)** Screenshot of SliceIt showing three different search options currently available in the database i.e. Search by gene name or Ensembl ID, Search by coordinates and Search by RBP name at various efficiency and specificity thresholds. **(C)** Figure describing various components of a typical search result page from SliceIt. Highlighted sections include Annotations, Data View, Exon expression with color coding and Genome browser view.

### 3.4 SliceIt as an integrative omics resource facilitates systematic experimental designs for editing RNA binding sites and their functional dissection across human tissues

SliceIt provides a robust set of sgRNAs that could be used for systematic perturbation of RBP’ binding sites occurring in a genomic loci or gene of interest. We have integrated several other omics datasets into SliceIt that can potentially help the users to define their criteria for RBP binding site centric Cas9 experiments. For instance – we investigated the gene PINK1 (PTEN induced putative kinase 1) with “search by gene” option in SliceIt. PINK1 is located at chr1:20,633,454-20,651,511. It is a forward oriented gene that encodes for three transcripts (longest isoform being the only annotated protein coding transcript). Pink1 also encodes for a long non coding RNA transcript (Pink1-AS) that is transcribed from this genomic loci (Figure 4A). SliceIt provides a total of 902 unique binding sites (499 binding sites in K562 and 665 binding sites in HepG2 cell lines with 21 common binding sites) in Pink1 genomic locus which are being targeted by 64 RBPs (based on eCLIP protein-RNA interaction profiles). SliceIt also provides a collection of 2667 unique sgRNAs (2345 sgRNAs in K562 and 2540 sgRNAs in HepG2 cell lines with 16 sgRNAs identified to target the sites in both the cell lines) around these binding sites. SliceIt also reports additional “omics” information such as SNPs, GWAS hits from the GWAS SNP catalog, tissue specific exon expression levels, distance from binding site etc. that can help the user for a site specific CRISPR/Cas9 experimental design. For instance – SliceIt provides 4123 SNPs reported in dbSNP database for PINK1 locus along with rs74315358 at chr1:20,644,549 locus from GWAS catalog. It also provides access to expression levels of exons so that the user can estimate a significant phenotypic difference around the binding site prior to employing Cas9 system for perturbation experiments. For instance, Fig. 4A shows the exon levels for PINK1 in brain frontal cortex generated by enabling the frontal cortex expression track, revealing that exon ENSE00001717360 is highly expressed.

**Figure 4.**
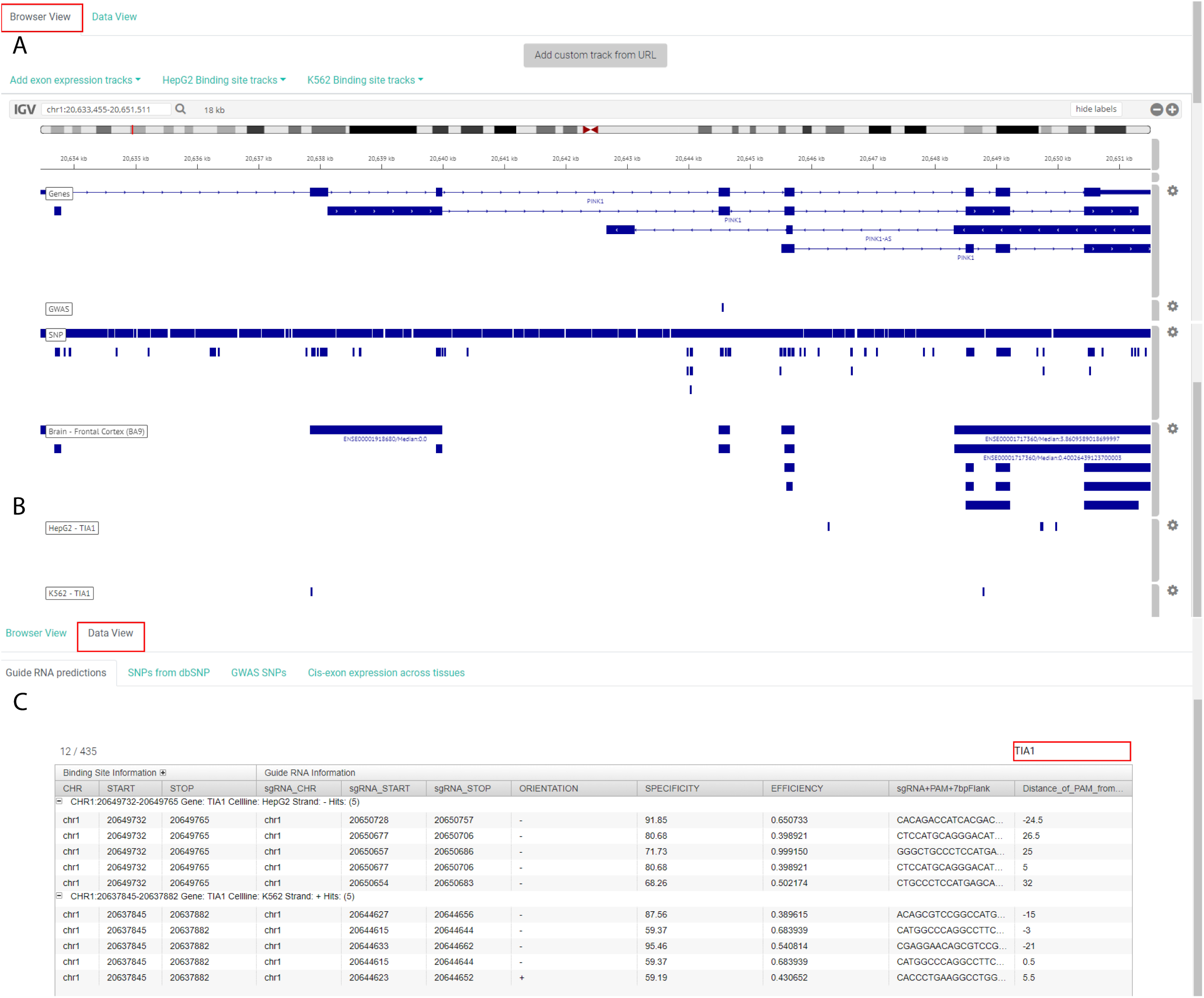
‘Browser View’ for Pink1 (PTEN induced putative kinase 1) queried using the “search by gene” option in SliceIt. (A) Panel showing multiple tracks for available transcripts of PINK1 along with their genomic coordinates, SNP tracks showing all SNPs from dbSNP database and GWAS hits occurring in this region, and exon expression track enabled for brain frontal cortex. (B) Panel showing the binding sites of TIA1 in both K562 and HepG2 cell lines for Pink1 locus. ‘Data View’ of SliceIt showing the top five potential sgRNAs with high specificity and efficiency for TIA1 binding sites which can be extracted by querying for “TIA1” using the available search box (highlighted in right corner).

PINK1 is an important serine/threonine-protein kinase that protects the cells from stress-induced mitochondrial dysfunction [50–53]. This gene has been implicated in several neurological disorders [54–56]. To design a Cas9 experiment to perturb the RBP binding sites in this gene locus, user can select an RBP of interest. For instance, several studies have shown that TIA1 (a cytotoxic granule associated RNA binding protein) is involved in stress induced apoptosis [57, 58]. To further understand the functional association in between these two key stress responsive genes (i.e. PINK1 and TIA1), user can employ Cas9 system to investigate the mechanism at high resolution scale by perturbing the TIA1 binding site/s on PINK1. User can enable the binding site track for quick navigation of all binding sites of TIA1 in SliceIt ‘browser view’ as illustrated in Figure 4B. Since, SliceIt has a huge collection of sgRNAs, to narrow down the sgRNA list, we provide sgRNAs predicted for respective binding site of an RBP in the ‘Data View’. For instance, in ‘Data View’, user can extract top 5 potential sgRNAs with high specificity and efficiency for each binding site, simply by querying for “TIA1” (Figure 4C). However, SliceIt provides a parametric flexibility for end users to choose sgRNAs based on their experimental criteria.

### 3.5 SliceIt aids in functional validation of RBP binding sites using CRISPR/Cas9 experiments

In order to validate the designed sgRNAs reported in SliceIt, we employed the “cause” and “effect” model of regulation by perturbing equivalent binding site on DNA using Cas9 genome editing system and study the impact of the perturbation on proximal exon expression levels (Figure 5A). Since SliceIt provides a user friendly platform to customize the choice of sgRNA selection based on user defined criteria, we used it to extract the list of sgRNAs targeting each binding site under consideration (with ± 50 bp flank sequence). We selected the sgRNAs which have efficiency > 0.70 and specificity > 70 % with minimal distance between on-site PAM (Protospacer Adjacent Motif) and center of the binding site. We also navigated the binding sites in ‘Browser View’ of SliceIt to verify if the sites are accompanied by at least one GWAS or dbSNP annotations. SliceIt also enables the exon level expression profiles. Hence, we also confirmed if the exon proximal to the binding site is expressed > 1 FPKM in primary tissue samples from GTEx project, corresponding to the cell line model being used. For this particular study, we used SliceIt to design potential sgRNAs for perturbing the binding sites and selected two sgRNAs for editing two different binding sites on LIPT1 (Lipoyltransferase 1). LIPT1 encodes for an acyl group transferase, involved in lipoic acid metabolism [59, 60] and glycine degradation [61]. Mutations in human lipoyltransferase gene LIPT1 were found to cause Leigh disease as well as Lipoic acid biosynthesis defects [59–61]. We navigated LIPT1 genomic loci and queried for two RBPs independently for respective binding sites (and predicted sgRNAs) as shown in Figure 5B (see Materials and Methods). We focused on the second exon of LIPT1 and investigated the impact of RBP binding sites (BS1 and BS2) targeted by respective sgRNAs (Lg1 and Lg2) designed by SliceIt (Figure 5B). We confirmed LIPT1 to be significantly expressed in primary liver tissue samples, since our cell line model is HepG2 cell line. Our main objective was to validate if these sgRNAs are likely to perturb the binding sites in HepG2 cells, with a high efficiency as predicted by SliceIt. Perturbation can result in increase or decrease in proximal exon expression levels, since it depends on whether the binding site can enhance or repress the activity of the exon i.e, binding site can be a splicing enhancer or repressor. SgRNA plasmid library constructs were confirmed using Sanger sequencing (Supplementary text 1). Plasmid transfection and lentiviral transduction efficiency was measured as GFP signals on fluorescence microscopy. GFP signals detected, confirmed 80-100% transfection efficiency in HEK293T cells and 75-90% transduction efficiency in HepG2 cells. For gene LIPT1, primers were designed to estimate the abundance of the second exon to validate the effect of two proximal binding sites’ (i.e. chr2:99157353-99157403 and chr2:99159478-99159514) perturbations using two different sgRNAs (labeled as Lg1 and Lg2 in Figure 5B) (see Materials and Methods). Upon normalizing with housekeeping gene PPIA, qPCR results showed significant (p<0.001) decrease in the exon expression levels compared to wild type HepG2 exon expression levels once transduced with Lg1 and Lg2 lentiviruses, respectively (Figure 5C). Hence, qPCR results confirmed that the designed sgRNAs targeted the binding sites, resulting in significant reduction of the proximal exon expression levels as a result of the perturbation of the binding sites. Interestingly, sgRNA – Lg2, designed to target the distal binding site of exon 2 exhibited higher reduction of the exon level than the sgRNA – Lg1 designed to perturb the proximal binding site.

**Figure 5.**
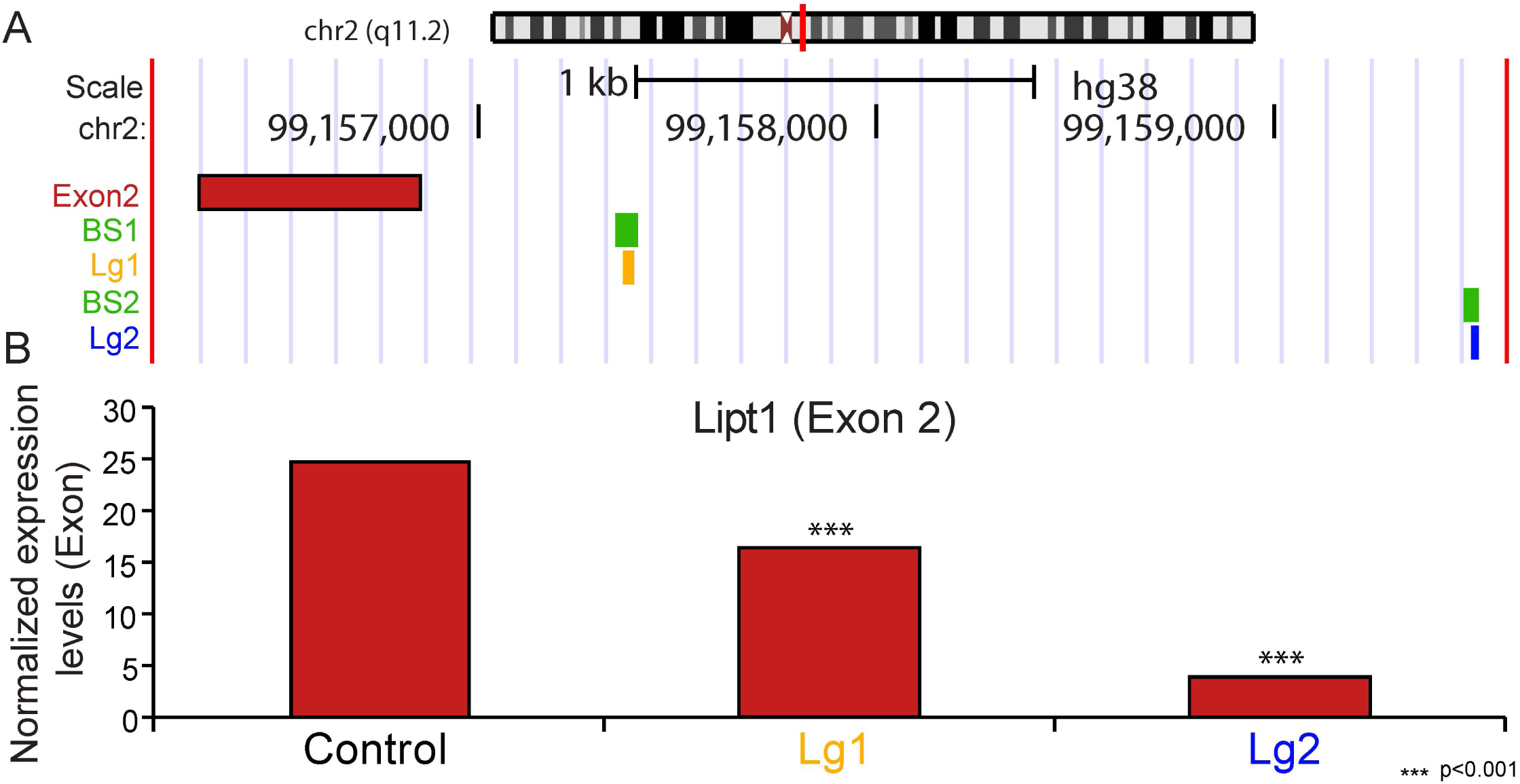
Functional validation of RBP binding sites using CRISPR/Cas9 experiments in human cell lines. **(A)** A hypothesis driven CRISPR/Cas9 model for determining the effect of sgRNA on an exon proximal to targeted binding site **(B)** A genome browser view of SliceIt illustrating the genomic loci of LIPT1 gene, the second exon, RBP centric query for binding sites (BS1 and BS2 color coded boxes) targeted by respective sgRNAs (Lg1 and Lg2, color coded boxes) designed by SliceIt. Since several sgRNAs can be predicted for a given binding site and multiple binding sites can be proximal, sgRNA track might appear dense and overlapping. **(C)** Binding site perturbation experiments using sgRNAs; Lg1 and Lg2. Exon expression levels (normalized) proximal to binding sites were quantified by qPCR (*** = p<0.001).

## 4. Conclusion

SliceIt is a comprehensive resource and visualization platform that enables the users to systematically design experiments to study the impact of a binding site of RBP on a particular RNA target. Hence, it can help the users in dissecting the role of binding sites in A) modulating splicing, stability and localization of RNA B) controlling the protein isoform levels, across a multitude of tissue types and cell lines by facilitating the generation of high quality custom set of sgRNAs for the well-established CRIPSR/Cas9 genome editing system. It is a one-stop repertoire that enables the design of small scale experiments to study a specific binding site’s role in modulating post-transcriptional regulation or medium range studies such as RBP centric functional screens or genome-scale CRISPR/Cas9 screens to edit the protein-RNA interaction networks. SliceIt also enhances the applicability of CRISPR/Cas9 system by focusing on binding sites on lncRNAs that can be perturbed for studying their contribution in downstream post-transcriptional control. Additionally, the “custom track” feature of SliceIt enables the users to re-purpose the compendium of SliceIt according to their choice. For instance, users can modulate the regulome for a novel RBP of interest and study the results in the context of existing protein-RNA interaction maps and expression profiles across tissue types.

## Author Contributions

SV, RS, and SCJ conceived and designed the study. RS and SH implemented the bioinformatics tool and integrated the datasets. SV, RS and SCJ designed the database. RS, SV and QM performed data analysis. QM, RS, XCD and SCJ designed the strategy to validate the predicted sgRNAs experimentally. SV, RS, QM, XCD and SCJ interpreted the data and wrote the manuscript. All authors read and approved the final manuscript.

## Acknowledgement

This work was supported by the National Institute of General Medical Sciences of the National Institutes of Health under Award Number R01GM123314 (SCJ).

